# FDJD: RNA-Seq Based Fusion Transcript Detection Using Jaccard Distance

**DOI:** 10.1101/2021.11.17.469019

**Authors:** Hamidreza Mohebbi, Nurit Haspel

## Abstract

Gene fusions events, which are the result of two genes fused together to create a hybrid gene, were first described in cancer cells in the early 1980s. These events are relatively common in many cancers including prostate, lymphoid, soft tissue, and breast. Recent advances in next-generation sequencing (NGS) provide a high volume of genomic data, including cancer genomes. The detection of possible gene fusions requires fast and accurate methods. However, current methods suffer from inefficiency, lack of sufficient accuracy, and a high false-positive rate. We present an RNA-Seq fusion detection method that uses dimensionality reduction and parallel computing to speed up the computation. We convert the RNA categorical space into a compact binary array called *binary fingerprints*, which enables us to reduce the memory usage and increase efficiency. The search and detection of fusion candidates are done using the Jaccard distance. The detection of candidates is followed by refinement. We benchmarked our fusion prediction accuracy using both simulated and genuine RNA-Seq datasets. Paired-end Illumina RNA-Seq genuine data were obtained from 60 publicly available cancer cell line data sets. The results are compared against the state-of-the-art-methods such as STAR-Fusion, InFusion, and TopHat-Fusion. Our results show that FDJD exhibits superior accuracy compared to popular alternative fusion detection methods. We achieved 90% accuracy on simulated fusion transcript inputs, which is the highest among the compared methods while maintaining comparable run time.

## Introduction

A *fusion gene* is a hybrid gene which is the result of two genes combined together. Gene fusion events are relatively common in many cancers including prostate, lymphoid, soft tissue, breast, and lung [1]. An increasing number of genes involved in infusions have been found to be promiscuous in that they can recombine with many different partner genes [2]. Gene fusions events often result from genomic structural rearrangements, transcription read-through, or abnormal RNA splicing [3, 4]. Two different transcripts from distinct genes may produce trans-splicing, which creates fusion transcripts. This trans-splicing is often a part of the normal RNA processsing [5] and is rather common among lower eukaryotes [6, 7]. Additionally, fusions may be produced by adjacent genes yielding a single, joined RNA product. This can create a read-through transcript or a conjoined gene [8]. Perhaps one of the best known examples of this type of fusion event is **BCR-ABL1** [9], which is the product of chromosomal translocation and is found in Leukemia patients. For this reason, studying gene fusion events is important for our understanding of cancer and for developing treatment options.

DNA and RNA high-throughput sequencing methods provide an opportunity to understand and characterize the transcriptome [10]. It is also useful in identifying fusion gene fusion events. Whole Genome Sequencing (WGS) is a technique for analyzing an entire genome and possibly finding abnormalities such as gene fusions. *RNA-Seq* [11] is a technology that uses Next-Generation Sequencing (NGS) to reveal the presence and quality of RNA in a biological sample. Transcriptome sequencing by RNA-Seq represents the “expressed exome” of the tumor itself and is highly suited for fusion transcript detection. This means, RNA-Seq captures only the transcriptionally active regions of the genome that reflects any rearrangements, and therefore, it reduces the sequencing costs compared to WGS [12]. Maher et al. [13] demonstrated the potential of this technology by applying transcriptome sequencing to several examples including the *TMPRSS2-ERG* gene fusion [14].

We can classify the methods to identify fusion transcripts from DNA [4, 15] and RNA-Seq data [16, 17] into two categories: *mapping-first* and *assembly-first*. Mapping-first methods perform genome alignment to identify discordantly mapping reads suggestive of rearrangements. An assembly-first approach first assembles the input reads into longer transcript sequences. Then it tried to find chimeric transcripts consistent with chromosomal rearrangements. In order to find evidence of gene fusion events, we can count the number of RNA-Seq fragments found as split (junction) reads that have direct overlap with a chimeric junction. Another possible indication of a junction is a paired read where each read maps to opposite sides of the chimeric junction without directly overlapping with the junction itself. Different fusion detection methods use various read alignment tools, genome databases, and criteria for reporting candidate fusion transcripts and exclude false positives. This leads to differences in accuracy, installation complexity, execution time, robustness, and hardware requirements. With the increasing amounts of data, there is a need to develop a fast and accurate fusion detection method. Current methods process samples that may contain up to tens of millions of reads. Processing these files takes up to days and the resulting output contains hundreds to thousands of gene fusion candidates, most of which are likely false or have little supporting evidence.

Whole-genome short-read sequencing is now routine and affordable. However, analyzing the genome remains a challenge due to the very large number of small, overlapping reads which cover the genome. This results in a true “Big Data” problem where we aim to find a small number of chromosomal breakpoints among tens of millions of reads. Efficient search and parallelization methods are required to carry out this task. In our work we present an efficient, parallel computing based method to reduce the dimensionality of the search space by splitting the candidate reads into windows and representing them using a compact binary format. We search for the location of possible gene fusions in the cancer genome using a Single instruction, Multiple Data (SIMD) parallel algorithm with the Jaccard distance as the similarity measure. The dimensionality reduction search technique takes advantage of an SSE multi-thread architecture to achieve efficient search, as well as the speed and accuracy of the STAR RNA-Seq aligner [18], a method capable of unbiased de novo detection of canonical junctions, as well as non-canonical splices and chimeric (fusion) transcripts.. In this work we focus on detecting non-contiguous fusions on the genome, which are made of two paired reads in which each read is partially aligned to different chromosomes. It should be noted that the algorithm can be readily extended to detect fusions of genes on the same chromosome. This is the subject of future work.

This manuscript is an extension of our conference publication [19], in which we presented the FDJD method. FDJD generates fusion candidates using both split reads and discordantly aligned pairs which are produced by the STAR alignment step.

The structure of this paper is as follows: Section 1 discusses the proposed FDJD fusion detection method. Section 2 provides the benchmarking results of FDJD using both simulated and cancer cell RNA-Seq data. Our method and state of the art fusion detection methods are compared using accuracy and time measures. The following outlines the main goals of this research:

- Propose a novel method to detect the exact locations of cancer fusion events using an exact comparison of given query with potential gene fusion breakpoint at reference genome.
- Provide an efficient data conversion method to reduce the dimensionality of data from categorical to binary formats.
- Provide an efficient parallel algorithm using both the multi-threaded environment and SSE instructions for exact Jaccard distance computation.
- Analyze the effectiveness of the proposed method on both simulated and genuine RNA-Seq data sets.

## 1 Fusion Detection using Jaccard Distance (FDJD) Method

### 1.1 Description

In our conference work that provides the basis for this extension, we described the Fusion Detection using the Jaccard Distance (FDJD) approach, which takes Illumina RNA-Seq data as input and generates lists of candidate fusion transcripts as output [19]. This pipeline comprises the alignment using the STAR aligner [12, 18], candidate generation, and refinement, detailed below. In this extended version we provide a more extensive comparison of runtime and accuracy. Our results and analysis are based on a larger number of examples.

Figure 1 shows an overview of the FDJD method, which includes the following parts. Part A) The FDJD fusion detection pipeline which comprises the alignment, candidate generation, and refinement steps. The FDJD method converts query sequences into a novel binary format, which then enables us to directly and efficiently search for fusions on the reference genome. Part B) The parallel Jaccard distance computation algorithm uses a tiling mechanism to load one block of data from each input into the cache memory. Part C) shows an example of the BCR-ABL1 gene fusion between chromosomes 9 and 22 [2]. The rest of this section details each step of the FDJD pipeline.

**Fig 1.** A) The FDJD fusion detection pipeline which comprises the alignment, candidate generation, and refinement steps. The FDJD method converts query sequences into a novel binary format, which then enables us to directly and efficiently search for fusions on the reference genome. B) The parallel Jaccard distance computation algorithm uses a tiling mechanism to load one block of data from each input into the cache memory. C) The BCR-ABL1 gene fusion between chromosomes 9 and 22 [2].

### 1.2 Alignment and Discordant Read Detection

The first task is the alignment step performed by the STAR alignment tool [12] that generates aligned BAM and chimeric output files. Following the alignment step, the discordant read detection phase takes the aligned pair-ended short reads as input. A discordant read pair is a pair that not aligned to the reference genome with the expected distance or orientation. If a set of discordant read pairs are mapped to two different genes, a fusion gene is suggested. Discordant read pairs are those that abnormally align to the reference; their mapping is somehow different than what is expected. A sequence read that spans a fusion event is called a splitting read. If both splitting parts of a read can be mapped and its mate is uniquely mapped to the reference genome, the splitting read is further marked as a discordant read-pairs. A sequence read can be considered as a fusion candidate when a cluster of overlapping discordant pairs satisfies two criteria: 1) there must be at least *minD* discordant read pairs in the cluster (Default value is 3), and 2) there must be at least *minS* soft-clipped reads from either side of the event that remap within the cluster region, which is the region from the outermost read in the cluster towards the variant breakpoint (Default value is 3, too). In this work we considered a read as a fusion candidate if pair mate positions were at least 500 base pairs apart from each other. Also, we only considered high quality reads with score greater than equal to 30.

### 1.3 Binary Fingerprint Generation

One of the main challenges of fusion detection is the huge search space. We need to compare hundreds of millions sample reads with billions of reference genome reads. One way to increase the efficiency of the search is reduce the dimensionality of the data and therefore reduce the data communication overhead. In this research, the inefficient categorical space is converted into a compact binary array, or *binary fingerprint*. The binary fingerprint format has previously demonstrated its effectiveness in the detection of structural variation breakpoints using DNA data [15]. The binary fingerprint generation phase takes a set of discordant read pairs as input, as described in subsection 1.2. The first step of the binary fingerprint generation is to split the pair-ended reads of length *l* into windows of length *w*. In this research, we consider 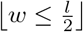 to be able to find breakpoints that are far from the middle of the read. Since in this work *l* = 50bp for our RNA-Seq read data, we considered *w* = 25. However, we used padding and set *w* = 32 in order to be able to load each binary fingerprint into one 128-bit SSE register for the next phase. We are taking advantage of the SIMD instructions [20] in the parallel Jaccard distance computation algorithm at the next phase 1.4. We convert each one of the query window strings of the then store its converted binary fingerprint format, detailed below.

Each window of length *w* is expressed as subsequences of length *k* or *k*-mers. First, all possible *k*-mers are placed in an indexed array. We call the indices of this *k*-mers array *fingerprints*. In other words, a fingerprint is just a sequence of indices in the array (e.g, [113442589]) representing the *k*-mers of a given window read. The following step is to convert a digit fingerprint array into a binary format. A binary fingerprint represents the index value associated with the existence of a *k*-mer of nucleotides in the sequence. That is, the base-10 indices array is converted into a binary indices array (e.g, [00110001010]). In this work, we considered *k* = 4. Since the input RNA sequence contains four possible letters (A, C, G, U – we skip N for simplicity), we have 4^4^ = 256 possible 4-mers. However, a *k*-mer and its reverse complement can be represented by the same index resulting in 128 possible indices. Therefore, each fingerprint contains 128-bit positions. The binary representation of a fingerprint uses one bit for each index in fingerprint – 0 if it is not represented in the window sequence, 1 otherwise. We are then able to load the binary fingerprint of each window into one SSE register and then perform SSE instructions on the reference genome. This, in turn, improves the computation time. The binary fingerprints of the input read pairs are compared with the reference chromosome via Jaccard distance in the following phase, described in Section 1.4, which takes advantage of both SIMD and the tiling mechanism to achieve the highest possible performance.

### 1.4 Parallel Jaccard Distance Computation

In addition to converting the discordant reads into a binary fingerprint format, we also need the reference genome to be represented in the same way for this step. We can convert the reference human genome into binary fingerprint only one time, and it can be reused in subsequent experiments. Then, we perform the search for fusion candidates using a fast parallel algorithm to calculate the similarity between the query and reference fingerprints using the Jaccard distance. This parallel algorithm performs a tiling mechanism using a block of threads to calculate the Jaccard distance between all the pairs of query and reference input. We carefully calculate the amount of cache that we have and load enough data to fill the cache. We used the OpenMP [21] library to provide the required threads and shared memory infrastructure. Each OpenMP thread performs the Jaccard distance calculations using SIMD instructions to achieve data level parallelism. Figure 1, part B illustrates the parallel Jaccard distance algorithm using a tiling mechanism. The first step is to read the binary fingerprints, where the query and reference genome inputs are then processed by a window of *t* threads to access *t* elements of each input. Each thread computes the Jaccard distance as follows. Let *A* and *B* be two sets. Their *Jaccard index* is the cardinality of their intersection divided by that of their union:

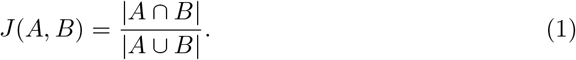

We can also rewrite Equation 1 into the following form. Equation 2 is conducive to faster computation than Equation 1.

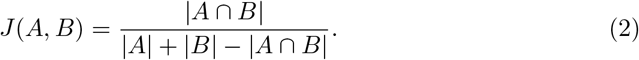

*J* (*A, B*) is between 0 and 1 and *J* (*A, B*) = 1 for the case that both *A* and *B* are empty sets. The *Jaccard distance* between *A* and *B* is 1 minus their Jaccard index:

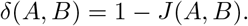

The input of this phase are two binary fingerprint queries with length of 128 bits. This enables us to load each query and reference data into a 128-bit SSE register [22] and perform the required SSE instructions including and/or and population count (*popcnt*) operations [23] to calculate *δ*(*A, B*). The popcnt operation counts the number of ones in a binary vector of length 128. This phase is implemented in C and uses *intrinsics* [24], which imposes SSE assembly instructions to the C program.

### 1.5 K-nearest Neighbors

The next task is to search the reference genome for the *K*-nearest neighbors of a given read query. The reads that are mapped to different chromosomes in the genome are considered as fusion candidates. The Jaccard distance results of the previous phase are processed at this phase and the most similar samples to a given query are selected. We take advantage of the observation that any system consisting of entities that interact pair-wise, such as the genes involved in fusions, can be described in terms of a network. Mitelman showed that 90% of the inter-connected networks of 358 known gene fusions in 2007 creates 3 clusters [2]. In our proposed method, we count how many times each query gene was involved in fusion with other genes and use it as a basis for a K-nearest neighbors (KNN) algorithm. We considered a range of values for *k* between 1 ≤ *k* ≤ 10; We found out that *k* = 3 achieves the best results that are aligned with the inter-connected network structure of gene fusions with 3 clusters [2, 25] The algorithm presented in this subsection is based on this idea that the *K*-nearest neighbors must be in a hypersphere with a radius determined by the triangle inequality [23]. Many researchers use GPUs for computing KNN in the Euclidean space [26]. Here, the task is to find indices of the *K* smallest values in a long array. Existing KNN methods are listed in chronological order: modified insertion sort [27], priority queue [28], probabilistic selection [29], and quicksort with different strategies for pivot selection [30]. Here we outline our solution:

Let *S* be a set of *n* binary vectors with length 128 and let *A* be a 128-bit binary fingerprint vector. The task is to find the maximum distance between each of the *K*-nearest neighbors of *A*. Let *B*_1_, *B*_2_, …, *B*_*K*_ be the *K*-nearest neighbors of *A*, sorted by their distances to *A*. We can set an upper bound for the distances among them using triangle inequality, for 1 ≤ *i* < *j* ≤ *K*:

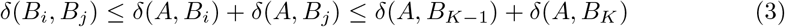

if *B_x_* is far from *A* such that *δ*(*A, B_x_*) > *δ*(*A, B*_*K*−1_) + 2*δ*(*A, B_K_*), then *B_x_* cannot be one of the *K*-nearest neighbors of *B_i_*:

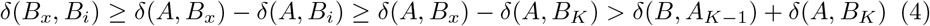

The *B_i_*’s already have at least *K* neighbors within this distance. This leads to the following algorithm to compute KNN: After finding the *K*-nearest neighbors of *A*, the method finds a subset 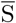 of *S* that contains vectors no farther than *δ*(*A, B*_*K*−1_) + 2*δ*(*A, B_K_*) to *A*. If 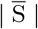 is less than | *S* | /2, we can find the *K*-nearest neighbors for *B*_1_, *B*_2_, …, *B_K_* with respect to 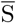. This algorithm works like truncated min-reduction, in which it repeatedly keeps the minimum of each pair of elements, and thus reduces the number of elements by half in each iteration. It stops the reduction when the next iteration will leave fewer than *K* elements. The pivot for partitioning is the maximum among the remaining elements. One partitioning is enough when *K* is small. The partition to the left of the pivot is simply sorted so that the *K* smallest distances and their indices are at right places. This pivot selection process is called *reduce-select* in the KNN algorithm 1. In order to find the best breakpoint location in a gene fusion candidate, we split it on several possible points. We then set a score based on how many times each gene was involved in fusions with other genes.

### 1.6 Refinement

Refinement is the last step. The input is both the chimeric disjoint and fusion candidates files. The initial list of candidates may include many false positives. Therefore, we conduct the refinement to further filter out unlikely candidates from the initial prediction. The refinement step minimizes the number of false positives by aligning the resulting reads against the human genome using BLAST sequence alignment [31]. In this work, a local BLAST database build was used in order to achieve higher speedup. The refinement phase considers only reads that are mapped to different chromosomes. The candidates are further grouped by gene pairing and breakpoint proximity. Then, they are further filtered by read support, extent of alignment at the putative breakpoint, and sequence similarity between putative fusion gene partners, performed in the following order:

1. **Grouping by breakpoint proximity.** We merge close fusion candidates into one query. Fusions with non-reference breakpoints that fall within a window of ±5 bases of each other are merged into a single fusion prediction.
2. **Assessing strength of alignment evidence.** Fusions that have only split-read support are required to have at least 25 bases of read alignment at each end of the breakpoint.
3. **Filtering sequence-similar fusion pairs.** We exclude fusion candidates where there exists a significant sequence alignment (BLAST, *E* ≤ 10^−3^) between any two reference isoforms of the genes.

#### Algorithm 1

The pseudocode of a new KNN algorithm using the reduce-select method.

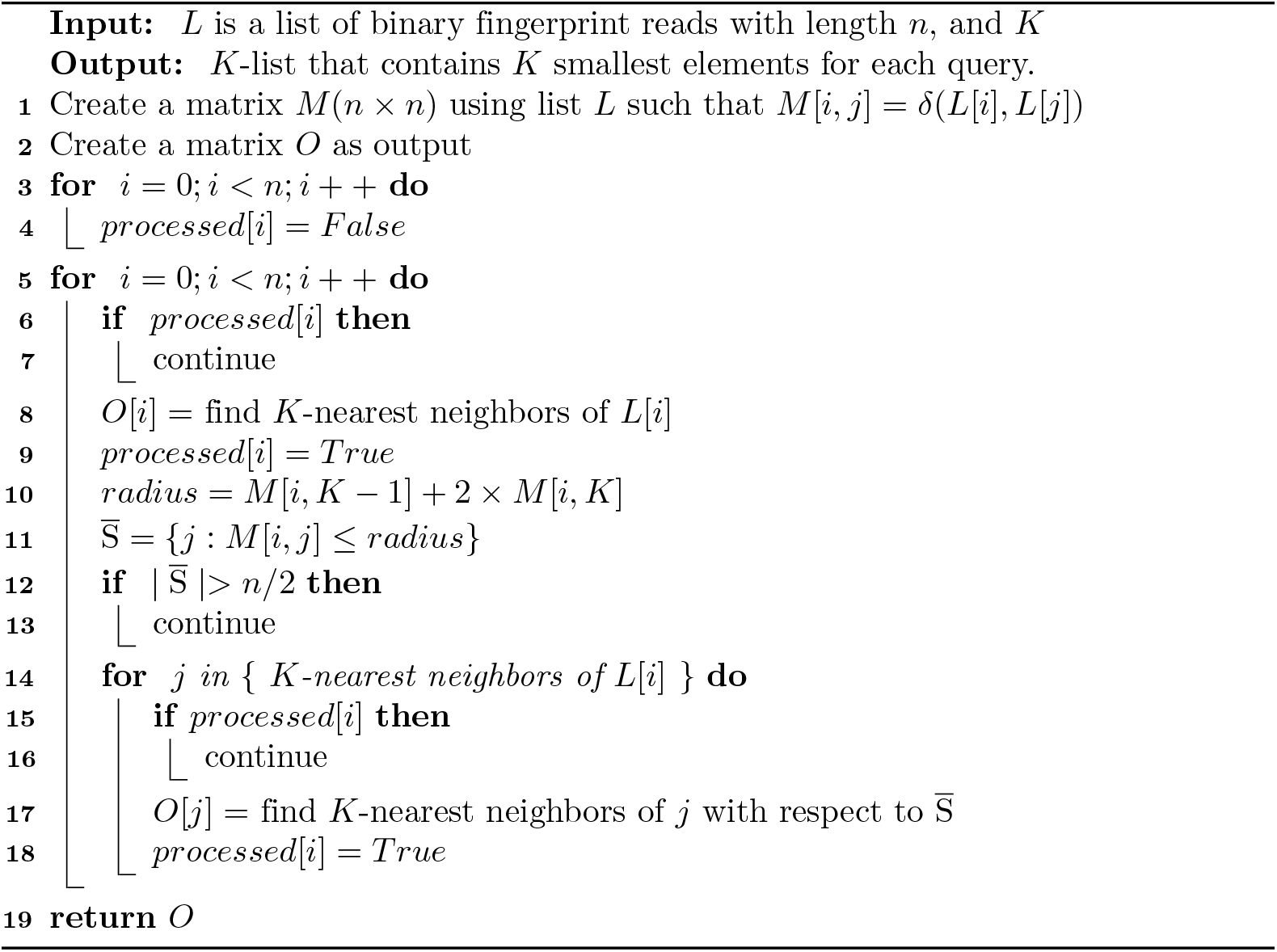

## 2 Results and Discussion

We used both simulated and genuine RNA-Seq datasets in our evaluations. Also, we compared the FDJD results with the following state-of-the-art fusion prediction methods: STAR-Fusion [12], Arriba [32], Pizzly [33], TopHat-Fusion [34], and InFusion [35]. The accuracy and run time comparisons are presented in Section 2.2.

### 2.1 Data Sets

The simulated dataset contains cancer cell lines generated by the *Fusion Simulator Toolkit* [36]. This simulator randomly selects two protein-coding genes from the Gencode v19 annotations. Then, it constructs a fusion transcript by randomly fusing a pair of exons randomly selected from each gene. This simulated gene fusion event requires that each gene contribute at least 100 bases of transcript sequence to the generated fusion. In our experiments, we made ten simulated samples, each containing 500 random fusions in 30 million PE Illumina RNA-Seq reads. Half of the simulated samples generated 50 base reads and the other half 100 base reads.

The fusion predictions of the studied methods analyzed genuine cancer line transcriptomes. The paired-end Illumina RNA-Seq data were obtained from 60 publicly available cancer cell line data sets, spanning a variety of cancer types including breast cancer (cell lines BT474, KPL4, MCF7, and SKBR3) and prostate cancer (cell line VCaP). To facilitate benchmarking and runtime analysis, 20 million paired-end reads were randomly sampled from each dataset and targeted for fusion prediction.

### 2.2 Fusion Identification Analysis

The accuracy evaluation of fusion identification results includes measuring the true positive (TP), false positive (FP), and false negative (FN) metrics. The challenge in benchmarking using genuine cancer cell RNA-Seq data is that the ground truth cannot be perfectly defined. Previously, an approach named *wisdom of crowds* [37] suggested that true fusions can be defined as those predicted by at least *n* = 3 different methods. We consider a false predictions as one that is predicted uniquely by any single method. We decided to consider non-unique fusions predicted by fewer than *n* different methods, as unsure (unscored) fusions. The TP and FP for fusion predictions were assessed at each minimum supporting evidence threshold to generate precision-recall curves. Additionally, we the measured prediction accuracy as the number of reads under the precision-recall curve (AUC). We believed that AUC is better suited than the popular receiver operating characteristic curve (ROC) for studies such as fusion prediction. This is because the number of true negatives (at least ~ 20*k*^2^, considering possible gene pairings) far exceeds the number of true positive fusions [38].

Figure 2 shows the AUC for the genuine data A) and simulated data B) for all the compared methods. This figure shows that FDJD achieved higher accuracy than the other methods. Part A) represents the AUC comparison of the studied fusion detection method for 92 breast cancer cell lines. These samples including four breast cancer cell lines *BT474*, *KPL4*, *MCF7*, and *SKBR3*. The FDJD method outperforms other method and achieves maximum AUC value of 0.9. The next best method is Arriba with 0.85 as its maximum AUC value. The mean of the AUC values also shows the better performance of the FDJD method. Part B) of this figure shows the AUC comparison for the simulated data. Every method except TopHat-Fusion performed well on the simulated datasets. Nearly all methods had improved accuracy with longer vs. shorter reads. FDJD outperformed the compared methods in combining chimeric and unmapped reads in addition to its increased accuracy in fusion detection.

**Fig 2.** The calculated area under the precision-recall curve (AUC) for FDJD and compared state-of-the-art fusion detection methods. Part A) represents the AUC value comparison of the studied method using genuine data. FDJD achieves the highest AUC values among studied methods. Part B) of this figure shows that the FDJD method surpassed other methods using simulated data with both 50 and 100 as base lengths (BL).

We evaluated the performance of all the methods on genuine datasets including four breast cancer cell lines: *BT474*, *KPL4*, *MCF7*, and *SKBR3*. We used the wisdom of crowds approach with *n* = 3 in building our ground truth dataset. Out of a total 86 fusions predicted by at least three methods, we found 44 fusions validated by at least three methods. We observed that as more methods predict the same fusions, the quality of validated fusions are improved via our definition of the fusion transcript truth sets. Therefore, rather than being limited to a single truth set, we could explore all possible truth sets defined by a range of values for *n* methods. We can then examine the distribution of leaderboard rankings for methods across all evaluated truth sets. We found out that the relative rankings were mostly stable regardless of which *n* was used to define the truth set. Table 1 compares the experimentally identified breast cancer fusions among the studied methods. This table shows that FDJD outperformed the other methods and was able to detect more validated fusions.

**Table 1.** This table shows the identification comparison of experimentally validated fusions in breast cancer cell lines *BT474*, *KPL4*, *MCF7* and *SKBR3*. All fusions identified by at least 3 different methods are shown. This table shows that FDJD outperformed the related methods and is able to detect more validated fusions.

|  |  | Fusion Names and Cancer Cell Lines |  |  |  |  |  |  |  |  |  |  |
| --- | --- | --- | --- | --- | --- | --- | --- | --- | --- | --- | --- | --- |
| Methods |  | EIF2AK1—PSMD3 | PPP1R3F—NUBPL | C5—FAM120A | CTC—340D7.1 | URM1—ATP8A2 | ADAMS19—SLC27A6 | BCAS4—BCAS3 | KIAA1919—SLC27A6 | NCOA2—CALB1 | SLCA5—SMG1 | TATDN1—GSDMB |
|  |  | BT474 | BT474 | KPL4 | KPL4 | KPL4 | MCF7 | MCF7 | MCF7 | SKBR3 | SKBR3 | SKBR3 |
|  | Arriba |  | ✓ | ✓ | ✓ | ✓ |  | ✓ | ✓ | ✓ |  | ✓ |
|  | Pizzly | ✓ |  | ✓ |  | ✓ |  | ✓ |  |  |  | ✓ |
|  | TopHat-Fusion | ✓ |  |  |  | ✓ | ✓ | ✓ |  |  |  | ✓ |
|  | STAR-Fusion |  | ✓ |  | ✓ | ✓ | ✓ | ✓ |  |  | ✓ | ✓ |
|  | InFusion | ✓ |  |  | ✓ | ✓ |  |  | ✓ | ✓ | ✓ |  |
|  | FDJD | ✓ | ✓ | ✓ |  | ✓ | ✓ | ✓ |  | ✓ | ✓ | ✓ |

Figure 3 illustrates the predicted fusions for the state-of-the-art methods in an UpSetR [39] style plot. This plot provides an efficient way to visualize the intersections of multiple sets compared to the traditional approaches like the Venn Diagram. Each bar and its value is compared with the number of circles associated with each of the compared methods. For example, the number 33 on the left side, with all methods checked at the bottom of the plot, indicates that 33 fusion samples were identified by all compared methods. The FDJD method identified the largest number of fusions and achieved the best ranking across methods in most cases. This ranking is followed by STAR-Fusion and Arriba. Notably, the top two ranked methods all leverage the STAR aligner, which shows the effectiveness of this alignment tool for fusion detection.

**Fig 3.** This figure illustrates experimentally validated fusions in breast cancer cell lines BT474, KPL4, MCF7 and SKBR3. All fusions are identified by at least three different methods are shown and compared with other methods in an UpSetR [39] style plot. Each bar and its value needs to compared with the number of circles associated to each state of the art methods compared. For example, the number 33 and all methods checked indicates that 33 fusion samples were identified by all compared methods.

### 2.3 Run Time Analysis

Figure 4 compares the speedup between the serial code and the parallel tiling implementation discussed in Section 1.4, In our experimental settings, we used 16 concurrent OpenMP threads to compute the Jaccard distance. The figure shows the speedup achieved by the parallel algorithm. Figure 4 (B) compares the execution time of five simulated samples (*S*1, *S*2, …, *S*5) with 100 as base length. It should be mentioned that the Jaccard distance calculation is only one of the steps of the proposed pipeline, and therefore the total execution time of the FDJD method is different from the time shown in the figure.

**Fig 4.** A) Distribution of execution times for both simulated and genuine cancer cell lines. FDJD adds a post-processing step to the common fusion detection methods which adds to execution time. Yet, it is still comparable to other methods. B) The speedup gained by the parallel tiling mechanism presented in phase 3 of the pipeline (Figure 1. In our experimental settings we used 16 concurrent OpenMP threads for the Jaccard distance computation phase. The speedup achieved by the parallel algorithm is calculated over 5 different simulated samples (*S*1, *S*2, …, *S*5) with 100 as read base length.

Figure 4 (A) depicts the execution time comparison between different methods. As seen, the computation time varied dramatically between methods. The fastest method is Pizzly which is an alignment-free *k*-mer-based approach, followed by Arriba and STAR-based methods such as STAR-Fusion and InFusion. The reason FDJD was not as fast as some of the other method is that it is not an alignment-free approach, and it also has a “post-alignment” step which contributes to the accuracy but consumes time. FDJD takes the output of an alignment and then compares candidates with the reference genome converted into a binary fingerprint format. Also, it uses both the chimeric junction alignment output and the input BAM file. This step has to be performed serially and therefore the runtime is affected. Despite that, FDJD’s execution time is still comparable with the other methods. It should be noted that the main goal in developing our method is to improve the accuracy over existing methods. Therefore, when taking into account both the accuracy and execution time, FDJD is not the fastest method, but it achieves comparable execution time and the highest accuracy among the compared methods.

### 2.4 Availability of Source Code

The Github repository of the FDJD pipeline contains the source code and instructions to run the code. It is available through the following link: https://github.com/mohebbihr/FDJD.

## 3 Conclusion

Identifying gene fusions is crucial for cancer treatment since these events can be potent drivers of tumorigenesis and appear in many different types of cancer. State-of-the-art fusion detectors suffer from a lack of accuracy and often have a high false-positive rate. In this research, we described a fusion detection algorithm using Jaccard distance (FDJD). Our method converts discordant reads into a binary fingerprint format to reduce the huge dimensionality of the search space. A parallel tiling search algorithm is used to compare query fingerprints with the RNA reference genome, and then locates fusion transcript candidates. The Jaccard distance is used as a similarity measure to compare the query and reference genomes. This enables us to find the nearest neighbors of a given query using a fast *K*-nearest neighbors algorithm. We used both simulated fusion dataset and genuine cancer cell RNA-Seq data in our evaluation. We compared our fusion detection results with state-of-the-art methods like STAR-Fusion, InFusion, and TopHat-Fusion. FDJD reached 90% accuracy on simulated data and outperformed other methods. The highest accuracy was also achieved using genuine cancer datasets. FDJD achieved the highest accuracy and while it was not the fastest method, the run time is comparable to other methods.

## Acknowledgements

We thank Dr. Ming Ouyang for his help. We thank Dr. Dan Simovici and Joyce Quach for their earlier contributions to the study.

## References

1. Kim H, Cho G, Han S, Shin J, Jeong E, Song S, et al. Novel fusion transcripts in human gastric cancer revealed by transcriptome analysis. Oncogene. 2014;33(47):5434.

2. Mitelman F, Johansson B, Mertens F. The impact of translocations and gene fusions on cancer causation. Nature Reviews Cancer. 2007;7(4):233–245.

3. Weckselblatt B, Rudd MK. Human structural variation: mechanisms of chromosome rearrangements. Trends in Genetics. 2015;31(10):587–599.

4. Dehghanpoor R, Ricks E, Hursh K, Gunderson S, Farhoodi R, Haspel N, et al. Predicting the effect of single and multiple mutations on protein structural stability. Molecules. 2018;23(2):251.

5. Rajkovic A, Davis RE, Simonsen JN, RoTTMAN FM. A spliced leader is present on a subset of mRNAs from the human parasite Schistosoma mansoni. Proceedings of the National Academy of Sciences. 1990;87(22):8879–8883.

6. Krause M, Hirsh D. A trans-spliced leader sequence on actin mRNA in C. elegans. Cell. 1987;49(6):753–761.

7. Sutton RE, Boothroyd JC. Evidence for trans splicing in trypanosomes. Cell. 1986;47(4):527–535.

8. Akiva P, Toporik A, Edelheit S, Peretz Y, Diber A, Shemesh R, et al. Transcription-mediated gene fusion in the human genome. Genome research. 2006;16(1):30–36.

9. Shtivelman E, Lifshitz B, Gale RP, Canaani E. Fused transcript of abl and bcr genes in chronic myelogenous leukaemia. Nature. 1985;315(6020):550.

10. Carninci P. Is sequencing enlightenment ending the dark age of the transcriptome? nature methods. 2009;6(10):711.

11. Wang Z, Gerstein M, Snyder M. RNA-Seq: a revolutionary tool for transcriptomics. Nature reviews genetics. 2009;10(1):57.

12. Haas B, Dobin A, Stransky N, Li B, Yang X, Tickle T, et al. STAR-Fusion: fast and accurate fusion transcript detection from RNA-Seq. BioRxiv. 2017; p. 120295.

13. Maher CA, Palanisamy N, Brenner JC, Cao X, Kalyana-Sundaram S, Luo S, et al. Chimeric transcript discovery by paired-end transcriptome sequencing. Proceedings of the National Academy of Sciences. 2009;106(30):12353–12358. doi:10.1073/pnas.0904720106.

14. Tomlins SA, Rhodes DR, Perner S, Dhanasekaran SM, Mehra R, Sun XW, et al. Recurrent fusion of TMPRSS2 and ETS transcription factor genes in prostate cancer. Science (New York, NY). 2005;310(5748):644–648. doi:10.1126/science.1117679.

15. Mohebbi H, Vajdi A, Haspel N, Simovici D. Detecting chromosomal structural variation using jaccard distance and parallel architecture. In: 2017 IEEE International Conference on Bioinformatics and Biomedicine (BIBM). IEEE; 2017. p. 1959–1964.

16. Latysheva NS, Babu MM. Discovering and understanding oncogenic gene fusions through data intensive computational approaches. Nucleic acids research. 2016;44(10):4487–4503.

17. Wang Q, Xia J, Jia P, Pao W, Zhao Z. Application of next generation sequencing to human gene fusion detection: computational tools, features and perspectives. Briefings in bioinformatics. 2012;14(4):506–519.

18. Dobin A, Davis CA, Schlesinger F, Drenkow J, Zaleski C, Jha S, et al. STAR: ultrafast universal RNA-seq aligner. Bioinformatics. 2013;29(1):15–21.

19. Mohebbi H, Quach J, Haspel N. Fusion Transcript Detection from RNA-Seq using Jaccard Distance. In: proc. of ACM-BCB (in HPC-BOD workshop); 2020.

20. Mohebbi H. Parallel SIMD CPU and GPU Implementations of Berlekamp–Massey Algorithm and Its Error Correction Application. International Journal of Parallel Programming. 2019;47(1):137–160.

21. Dagum L, Menon R. OpenMP: An industry-standard API for shared-memory programming. Computing in Science & Engineering. 1998;(1):46–55.

22. Lomont C. Introduction to intel advanced vector extensions. Intel White Paper. 2011; p. 1–21.

23. Ouyang M. KNN in the Jaccard space. In: 2016 IEEE High Performance Extreme Computing Conference (HPEC). IEEE; 2016. p. 1–7.

24. Stojanov A, Toskov I, Rompf T, Püschel M. SIMD intrinsics on managed language runtimes. In: Proceedings of the 2018 International Symposium on Code Generation and Optimization. ACM; 2018. p. 2–15.

25. Mitelman F, Johansson B, Mertens F. Fusion genes and rearranged genes as a linear function of chromosome aberrations in cancer. Nature genetics. 2004;36(4):331–334.

26. Mohebbi H, Mu Y, Ding W. Learning weighted distance metric from group level information and its parallel implementation. Applied Intelligence. 2017;46(1):180–196.

27. Garcia V, Debreuve E, Nielsen F, Barlaud M. K-nearest neighbor search: Fast GPU-based implementations and application to high-dimensional feature matching. In: 2010 IEEE International Conference on Image Processing. IEEE; 2010. p. 3757–3760.

28. Barrientos RJ, Gómez JI, Tenllado C, Matias MP, Marin M. kNN query processing in metric spaces using GPUs. In: European Conference on Parallel Processing. Springer; 2011. p. 380–392.

29. Monroe L, Wendelberger J, Michalak S. Randomized selection on the GPU. In: Proceedings of the ACM SIGGRAPH Symposium on High Performance Graphics. ACM; 2011. p. 89–98.

30. Komarov I, Dashti A, D’Souza RM. Fast k-NNG construction with GPU-based quick multi-select. PloS one. 2014;9(5):e92409.

31. Kent WJ. BLAT – the BLAST-like alignment tool. Genome research. 2002;12(4):656–664.

32. Uhrig S, Fröhlich M, Hutter B, Brors B. PO-400 Arriba–fast and accurate gene fusion detection from RNA-seq data; 2018.

33. Melsted P, Hateley S, Joseph IC, Pimentel H, Bray NL, Pachter L. Fusion detection and quantification by pseudoalignment. BioRxiv. 2017; p. 166322.

34. Kim D, Salzberg SL. TopHat-Fusion: an algorithm for discovery of novel fusion transcripts. Genome biology. 2011;12(8):R72.

35. Okonechnikov K, Imai-Matsushima A, Paul L, Seitz A, Meyer TF, Garcia-Alcalde F. InFusion: advancing discovery of fusion genes and chimeric transcripts from deep RNA-sequencing data. PloS one. 2016;11(12):e0167417.

36. Fusion Simulation Toolkit;. https://FusionSimulatorToolkit.github.io.

37. Surowiecki J. The wisdom of crowds: Why the many are smarter than the few and how collective wisdom shapes business. Economies, Societies and Nations. 2004;296.

38. Davis J, Goadrich M. The relationship between Precision-Recall and ROC curves. In: Proceedings of the 23rd international conference on Machine learning. ACM; 2006. p. 233–240.

39. Conway JR, Lex A, Gehlenborg N. UpSetR: an R package for the visualization of intersecting sets and their properties. Bioinformatics. 2017;33(18):2938–2940.

